# HLAProphet: Personalized allele-level quantification of the HLA proteins

**DOI:** 10.1101/2023.01.29.526142

**Authors:** Michael B. Mumphrey, Ginny Xiaohe Li, Noshad Hosseini, Alexey Nesvizhskii, Marcin Cieslik

## Abstract

Loss of HLA expression in tumor cells is a commonly observed phenotype that is known to be associated with T-cell evasion. Proteogenomic characterizations of the molecular mechanisms underpinning this loss of HLA expression are hindered by the polymorphic nature of the HLA proteins, with most individuals having germline HLA sequences that are highly divergent from the sequences found in standard reference databases. To address this issue, we have developed HLAProphet, an algorithm that utilizes HLA types from paired DNA sequencing data to provide personalized allele-level quantification of the HLA proteins from TMT mass spectrometry data. We show that HLAProphet triples the number of tryptic peptide identifications made by standard reference based approaches, and produces protein expression values that have high concordance with RNA expression and known loss of heterozygosity events.

## 1. Main

It is now recognized that one of the body’s primary defenses against cancer is T-cell immunity^1–3^. To avoid this immune surveillance, malignant cells often develop phenotypes that disrupt T-cell activity, including loss of HLA function^4^. Proteogenomic characterizations of the HLAs are necessary to understand the molecular mechanisms underpinning loss of HLA function, however this is hindered by the highly polymorphic nature of the HLAs. For each HLA gene there are thousands of known alleles coding for thousands of unique proteins^5^. This is a problem for most molecular methods which rely on a standard reference containing a single sequence for each known gene. For the average individual, their genome will contain HLA sequences that are divergent from whichever sequences are present in the standard reference. For this reason, personalized approaches to genomic and transcriptomic analysis of the HLA genes have been developed^6^. Personalized proteomics methods for HLA quantification, however, are lacking.

A number of methods have been developed to allow identification and/or quantification of proteins containing variant sequences^7–12^. The general approach involves identification of germline or somatic variants using paired DNA sequencing data, *in silico* translation of these variant proteins, and then concatenation of these variant sequences with a standard protein reference to create an augmented search database. Unfortunately, these methods often restrict their analyses to single amino acid variants (SAAVs), which is not useful for the HLA proteins where we frequently observe multiple variants per tryptic peptide. Haplosaurus^13^ improved upon SAAV approaches by allowing quantification of proteins with an arbitrary number of amino acid variants by imputing complete phased genotypes into all protein coding genes before *in silico* translation. However, this generalized approach requires high quality genotypes. This is a problem for the HLA genes where their polymorphism precludes traditional genotyping, and where germline sequencing instead requires the use of specialized HLA typing software. This issue can be seen when looking at the 709 personalized HLA-A protein haplotypes inferred by Haplosaurus from 2504 samples from 1000 Genomes. Even though all 1000 Genomes samples have HLA types matching entries in the IMGT/HLA database^14^, none of the personalized HLA-A proteins reported by Haplosaurus can be found in the database [IMGT/HLA release 3.51.0]. This is likely caused by the presence of at least one error in genotyping or phasing in each sample, leading to the *in silico* translation of non-existent proteins. This demonstrates that solutions for personalized HLA quantification require a specialized approach that works from full HLA types, not individual variant calls. To address this we have developed HLAProphet, an algorithm that leverages the FragPipe software suite and the TMT-integrator^15^ algorithm to provide personalized quantification of the HLA proteins from multiplexed TMT mass spectrometry data using known HLA types.

To demonstrate the issues faced by traditional proteomics methods when quantifying the HLAs, we used FragPipe and the standard Gencode v34^16^ protein database to quantify all proteins in 208 samples from the CPTAC lung squamous cell carcinoma cohort. First, due to the polymorphic nature of the HLA genes, most HLA proteins will have tryptic peptides not found in a monomorphic reference database. This is evidenced by the HLA proteins having a significantly lower fraction of predicted tryptic peptides identified than other proteins of similar size and abundance **(Figure 1A)**. Second, when peptides are identified, abundance calculations are often done using incorrect assumptions about the number of times each peptide is coded for in the genome **(Figure 1B)**. This is an issue given that the number of copies of a peptide in an HLA type directly correlates to peptide abundance **(Figure 1C).** The most common case (41% of identified peptides) is the identification of peptides that are not actually present in an individual’s genome based on their HLA type. This occurs in multiplexed samples where a non-zero intensity value will be assigned to all aliquots in a plex, even if the abundance is actually zero, due to noise within the mass spectrometer. We can see this visually within plexes where peptides have much higher intensities within samples that are predicted to code for the peptide, when compared to those that do not **(Figure 1D-E)**. In the standard approach, this leads to the assumption that the peptide was lowly expressed, when in reality it was never coded for and provides no information about HLA expression. The next most common issue (35% of identified peptides) is when an individual is heterozygous for two different alleles of an HLA gene, with many unique peptides being coded for by only a single chromosome. This clashes with the standard assumption that an identified peptide is coded for twice in a diploid human genome. When calculating gene-level abundances, these peptides will lead to under-estimates of abundance since they only provide information for one of the two alleles for that gene. Finally, for the MHC class 1 genes, some peptides within the relatively conserved backbone can be shared across genes, but in a way that is not captured in the standard reference database sequences. This leads to the assumption that a peptide is unique to an HLA gene, when in fact there are more copies present than expected. For these peptides (5% of identified peptides), abundance will be overestimated. Only 19% of peptides meet the standard assumption of being diploid and unique to a single HLA gene. Finally, standard methods tend to report a single abundance for each gene. However, for each HLA gene, heterozygous haplotypes will generally code for two unique proteins, each with a unique peptide presentation repertoire. Therefore, it is important for a personalized HLA quantification algorithm to report allele level protein abundances. To address these issues we have developed HLAProphet, an algorithm that provides personalized allele level quantification of the HLA proteins.

**Figure 1.**
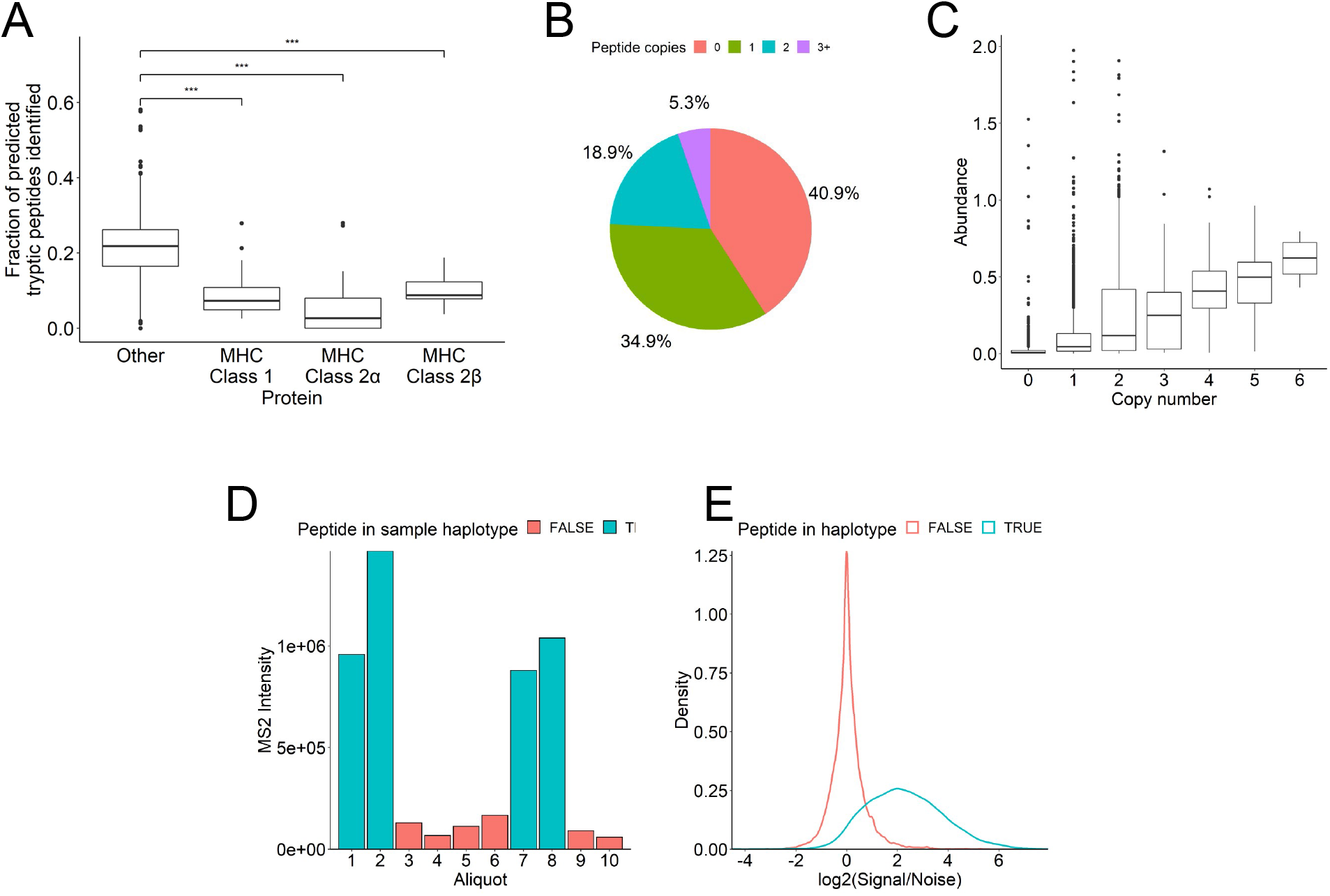
Issues faced when using standard reference (GENCODE v34) based proteomics to quantify HLA proteins in the CPTAC LSCC cohort (n = 208) using TMT mass spectrometry **(A)** Fraction of predicted tryptic peptides identified when analyzing tandem mass spectrometry data using a standard reference approach. Class 1, class 2α, and class 2β HLA proteins all have significantly reduced peptide identifications when compared to all other proteins of similar length and abundance. **(B)** True copy number of HLA tryptic peptides that are expected to be unique and diploid (exactly 2 copies) based on standard protein reference database GENCODE v34. **(C)** Correlation between abundance and true copy number for HLA tryptic peptides that are expected to be unique and diploid using a standard reference database. **(D)** Example MS2 intensities for a single peptide measured in a 10-plex. Each aliquot is colored based on whether or not the peptide is predicted to be coded for in that individual’s genome based on their HLA type. **(E)** Comparison of signal to noise ratios for 1,845 peptide identifications across the entire LSCC cohort. Noise is calculated as median MS2 intensity of aliquots that are not predicted to express a given peptide (red bars in E).

We next applied HLAProphet to the same set of samples to demonstrate the improvements in HLA peptide identification and protein quantification. HLAProphet uses HLA types derived from paired DNA sequencing data to generate a personalized HLA protein reference sequence database via *in silico* translation. We use here HLA types called by Hapster^17^, but any source of HLA types can be used. The personalized HLA database is then concatenated with an existing standard reference database, here GENCODE v34^16^, and is run through the FragPipe quantification workflow. The use of HLAProphet’s personalized HLA reference increases peptide identification by ~3-fold **(Figure 2A)**. Peptide spectrum matches are then used to quantify protein abundance using a personalized version of the TMT-integrator algorithm, which includes two major modifications. First, the standard approach assumes that all reference peptides are coded for in the genomes of all samples, which is not true for the HLA proteins. We therefore calculate HLA protein abundance on an individual basis, only using intensity values from peptides predicted to be present in a sample based on its HLA haplotype. Second, normalization between plexes is generally done by taking the ratio of a sample’s intensity to a pooled reference sample. The pooled reference acts as a physical average for tryptic peptide abundance in the standard case, given that all samples have the peptide coded for in their genome. However, for the HLA proteins, some peptides will only be coded for in a handful of samples, and therefore their average abundance in a pool of the entire cohort will be diluted by the samples with HLA types that do not code for the peptide. These diluted reference intensities shrink the denominator of the ratio calculation, causing inflated ratios and overestimates of peptide abundance in rare peptides **(Figure 2B)**. To address this issue, we multiply our reference ratios by a dilution factor **(Methods)**, removing the association between cohort population frequency and peptide ratio **(Figure 2C)**.

**Figure 2.**
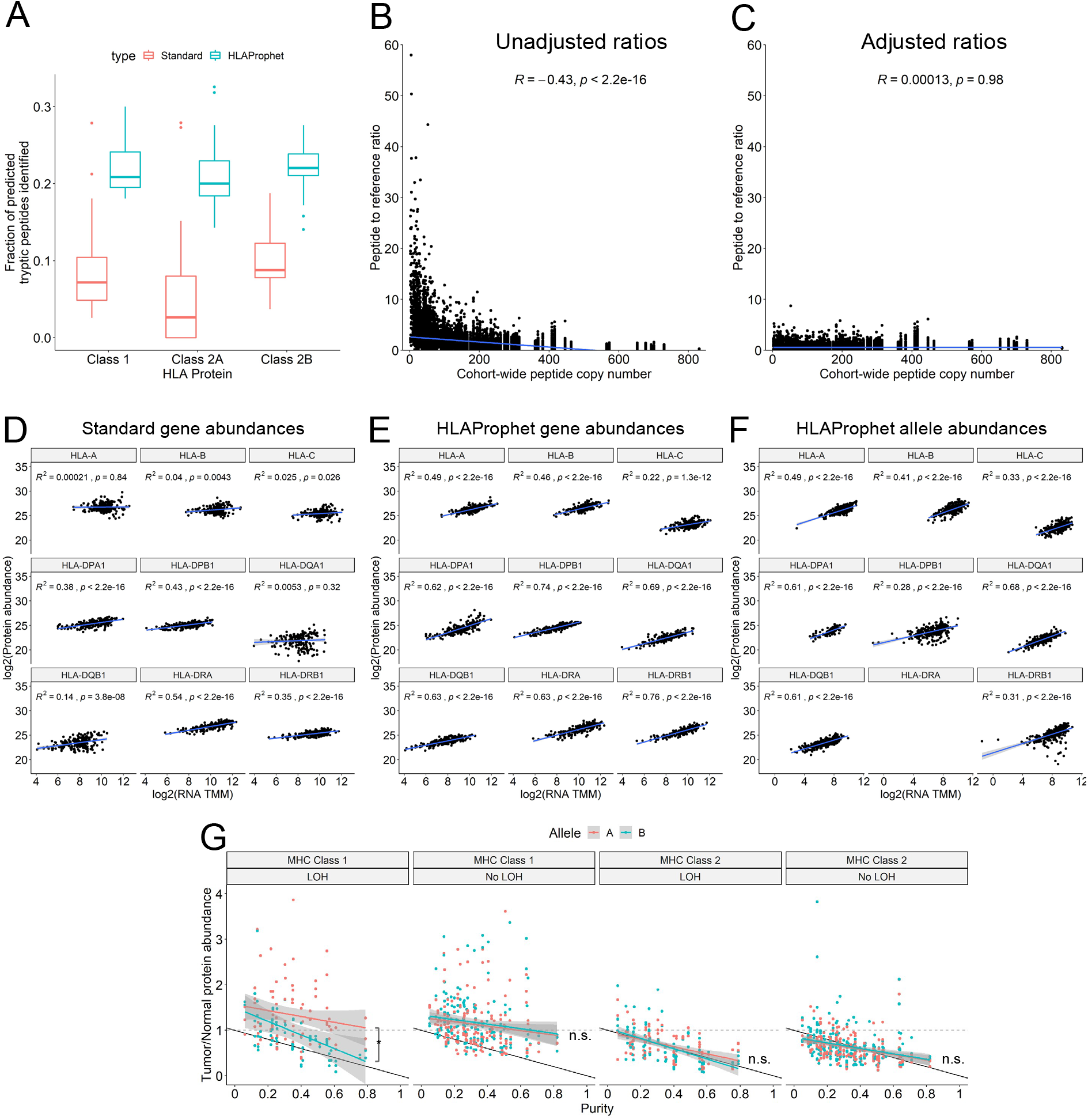
**(A)** Fraction of predicted tryptic peptides for HLA proteins identified using either the standard reference approach or HLAProphet. HLAProphet produces ~3x as many peptide identifications. **(B)** Ratio of peptide intensities to reference channel intensities for all identified HLA tryptic peptides compared to cohort-wide peptide copy number for the CPTAC LSCC cohort (n = 208). A standard unique diploid peptide would be found twice per sample for 416 total copies. HLA peptides have variable cohort representation, and rare peptides show a bias towards elevated ratios due to dilution in the reference channel. **(C)** Ratio of peptide intensities to reference channel intensities, adjusted for reference dilution. The bias caused by cohort copy number is completely removed. **(D-F)** Correlation of RNA expression to protein expression using a standard reference based approach **(D)**, or HLAProphet’s personalized quantification of gene level **(E)** or allele level **(F)** abundances for the CPTAC LSCC cohort. HLA-DRA is excluded from the allele level quantifications due to its low polymorphism, and low availability of allele-specific peptides. **(G)** HLA protein abundance in tumors as a fraction of normal tissue abundance. The “B” allele is the allele that is lost following an LOH event, while the “A” allele is the retained allele. Allele labels are randomly assigned for cases with no observed LOH events. Black lines show expected expression as a function of tumor purity, assuming the tumor cells have no expression. Significant differences between regression coefficients were tested using the Chow test.

To evaluate HLAProphet’s HLA protein quantification, we compared protein abundance to paired RNA expression for all samples. We see that when using the standard reference based approach, the MHC class 1 genes show no significant correlation between protein and RNA expression **(Figure 2D)**. For the MHC class 2 genes, most show significant correlation between RNA and protein expression, but with R^2^ values of only .14-.54. HLAProphet’s protein abundances improve correlations to RNA in all cases, with the MHC class 1 genes showing significant correlations and the MHC class 2 R^2^ values improving to .62-.78 **(Figure 2E)**. We also see that at the allele level, protein expression remains highly correlated with RNA expression for all genes, demonstrating HLAProphet’s ability to report allele level HLA protein abundances **(Figure 2F)**. We do, however, see a drop in R^2^ for HLA-DPB1 and HLA-DRB1. Interestingly, for HLA-DRB1, this loss of correlation is almost completely due to a single allele HLA-DRB*12:17 **(Supplementary figure 1)**, which shows no correlation between RNA and protein expression. It will take further experiments to determine if this is an artifact or a real allele-specific effect. HLA-DRA has low polymorphism and does not contain sufficient allele-specific peptides to quantify at the allele level.

To provide further evidence for HLAProphet’s ability to report allele level expression, we examined allelic HLA expression in tumor samples with loss of heterozygosity (LOH) events indicated by paired DNA sequencing data **(Figure 2G)**. We see that for the MHC class 1 proteins, in tumors with LOH the lost allele has significantly lower expression than the retained allele, with an effect size that increases with tumor purity. In tumors with no observed LOH, there is no clear difference between the two alleles. For the MHC class 2 proteins, which are not expected to be expressed in most tumor cells, we see no effect of LOH. However, we do see that the bulk HLA protein expression decreases exactly as expected with tumor purity under the assumption that tumor cells do not express the proteins.

In summary, we present here HLAProphet, a tool that enables personalized allele level quantification of the HLA proteins. We show that HLAProphet improves upon the existing state of the art standard reference based approaches by significantly increasing peptide identifications, and by providing protein expression values that show higher correlation to RNA and allelic loss of expression where predicted due to LOH. Moving forward, HLAProphet will enable the study of HLA loss at the protein level in tumors, providing greater insights into how cancers evade T-cell surveillance.

## 2. Methods

### Code availability

HLAProphet code is available at https://github.com/MBMumphrey/HLAProphet

### Data acquisition

Paired WES DNA sequencing, polyA RNA sequencing, and TMT mass spectrometry data were obtained for 108 paired tumor and normal samples from the CPTAC3 lung squamous cell carcinoma cohort.

### Standard reference based proteomics

Raw mzML files were run through the standard FragPipe quantification pipeline (https://fragpipe.nesvilab.org) for TMT mass spectrometry data. The GENCODE v34 protein database was used as the reference database for all peptide searches.

### HLA typing

HLA haplotypes were inferred using the Hapster software. For each sample, paired normal WES DNA sequencing data was used to predict HLA types.

### Tryptic peptide predictions

Tryptic peptide predictions for all proteins were performed using the cleave function from the python package Pyteomics. Predicted cleavages were performed for the trypsin protease, with minimum length 7 and maximum length 50, with up to 1 missed cleavage. For each HLA tryptic peptide, true copy number within a sample was determined by counting the number of times a tryptic peptide was predicted to be coded for in the HLA type of each individual sample.

### Personalized protein reference construction

HLA types for all aliquots are used to pull protein sequences from the IMGT/HLA database, which are then combined into a cohort-specific HLA fasta reference. For HLA types with different IDs, but with identical protein products, a single sequence is included in the final reference with a harmonized type ID. A relationship database is provided with a list of all original HLA types and their harmonized IDs, so that abundances can be tied back to their original HLA type during post-processing. The HLA reference database is then combined with a standard reference database (here GENCODE v34, with HLA sequences removed) using Philosopher, which produces reverse sequences and adds common contaminant sequences.

### Personalized protein abundance calculations

Protein abundances are calculated using the peptide spectrum match tables output by FragPipe, and the TMT-integrator algorithm (described at http://tmt-integrator.nesvilab.org/) with two major modifications. First, only peptides predicted to be coded for in a sample’s genome based on its HLA type are used, even if there are non-zero MS2 intensities reported for the peptide within that sample. Second, peptide to reference ratios are multiplied by a reference dilution factor, to account for dilution of rare peptides in the reference channel.

### Peptide ratio dilution factor calculation

To account for technical differences between MS/MS runs for an experiment with multiple multiplexed samples, a pooled reference sample is often included in one channel of each multiplex. Intensities can then be taken as a ratio to the reference channel to compare between plexes. This works well for most proteins where all uniquely mapping peptides are assumed to be coded for in the genome of all samples. In this case, the reference channel acts as a physical average of peptide expression across all samples in an experiment. For an experiment with N samples, the reference channel expression E is modeled as:

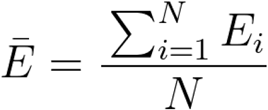

For the HLA proteins, many tryptic peptides will be coded for in only a subset of the sample genomes. However, the reference channel still physically averages all samples in the experiment, diluting the expression of a given peptide by combining it with other samples that cannot express it and therefore contribute no information about its expression. If we consider the experiment to be separated into P samples that code for the peptide and Q samples that do not code for the peptide, then we can assume the reference channel expression E is:

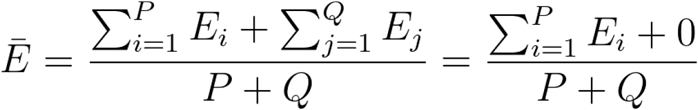

To remove the dilution effect of the Q samples that do not code for the peptide, we multiply by a dilution factor to produce the correct denominator for average abundance of P samples that code for the peptide:

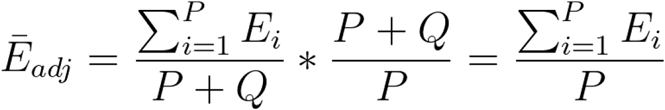

In practice, as P becomes very small, the dilution factor becomes very large, so we use a constant c to stabilize the value at the extremes for a final dilution factor:

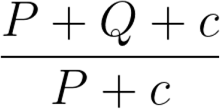

The ultimate goal is to produce peptide ratios that have no relationship with cohort population peptide frequency, so we test a range of c values to identify the value that best removes this relationship for use in the final calculations.

### MS2 signal/noise ratios

We assume that for peptides that are not coded for in an aliquot’s HLA type, the reported MS2 intensities are noise. Within each plex, we calculate the noise level as the median MS2 intensity of all aliquots that do not code for the peptide. We then take the ratio of all aliquot’s MS2 intensities to their plex and peptide specific noise value.

### HLA RNA expression quantification

Personalized genomic HLA reference sequences were produced for all samples using paired DNA sequencing data and the Hapster^17^ software. Genomic reference sequences were then spliced *in silico* to produce personalized transcript reference sequences. Allele specific RNA TPM values were calculated using Kallisto, personalized HLA transcript references, and polyA RNA seq data. For gene level TPMs, the sum of both HLA allele TPMs was used.

### Statistics

For the comparison of the predicted fraction of tryptic peptides identified, pairwise t-tests with Bonferroni corrections were performed. For correlations between peptide ratios vs cohort copy number, protein expression vs RNA expression, and tumor abundance vs tumor purity, linear regression was performed. To compare the regression line coefficients between A and B alleles for tumor abundance vs tumor purity, the Chow test was performed.

