## Supplementary figure 1 for "HLAProphet: Personalized allele-level quantification of the HLA proteins"

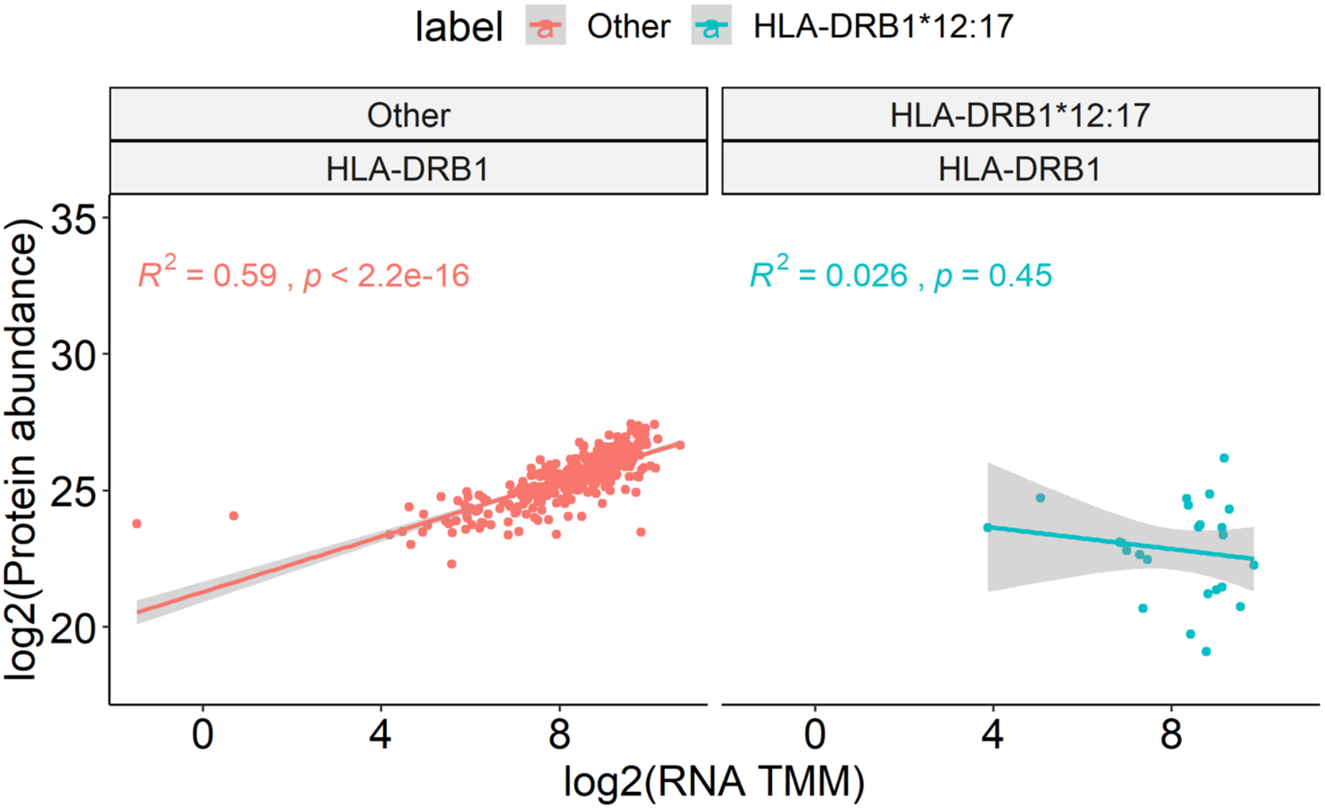


**Supplementary figure 1** - Correlation of allele specific RNA and protein expression for HLA-DRB1 (Data from Figure 2F) with HLA-DRB1*12:17 separated out. Most DRB1 alleles show good correlation between RNA and protein (left, red), with the exception of HLA-DRB1*12:17 (right, blue) showing no correlation.
